# Fine-tuned spatiotemporal dynamics of sporophylls in movement-assisted dichogamy: a study on *Clerodendrum infortunatum*

**DOI:** 10.1101/2020.05.30.124818

**Authors:** Amritendu Mukhopadhyay, Suhel Quader

## Abstract

Over 70% of flowering plants are hermaphroditic, with male and female parts in the same flower. Hermaphroditism is cost-effective because a common investment in reward and attractive structures yields benefits through both male and female reproductive success. However, the advantage is accompanied by an increased risk of self-pollen deposition, which is disadvantageous for both self-compatible and self-incompatible species. Hermaphroditic plants reduce self-pollen deposition by separating sporophylls (male and female reproductive parts) either spatially (herkogamy) or temporally (dichogamy). In movement-assisted dichogamy, both sporophylls are involved in a coordinated motion, where they move in opposite directions. However, the effectiveness of this adaptation in reducing self-pollen deposition may be compromised at the point when the sporophylls cross each other and are close enough to interfere, resulting in a transition phase problem. The solution to this problem lies in the details of the spatiotemporal dynamics of the sporophylls in relation to their reproductive maturity. We studied these details across the floral lifetime of a protandrous shrub *Clerodendrum infortunatum* (Lamiaceae), in rainforest fragments of the Western Ghats, India. We took photos of flowers at regular time intervals and measured sporophyll angles from the images. We also carried out a field experiment to determine stigma receptivity. The findings suggest that the effectiveness of dichogamy is maximised through two properties of the transition phase: physical resistance to self-pollen deposition by narrow stigma lobe opening, and chemical non-receptivity of the stigma during this phase. This study emphasises the importance of accessory adaptations in movement-assisted dichogamy to tackle the transition phase problem, which is inherent in this particular form of dichogamy.

## Introduction

Despite the existence of a diversity of mating strategies, over 70% of flowering plants are hermaphroditic (de Jong and Klinkhamer 2005). According to the theory of plant resource allocation in economic terms (Bazzaz and Grace 1997, Campbell 2000), hermaphroditism has a distinct advantage in reducing costs because the same accessory flower structures that provide rewards and advertisements are used to increase both male and female reproductive fitness (Charnov et al. 1976, Charlesworth and Charlesworth 1981, Barrett 2002). However, the associated costs are also high. In particular, hermaphroditism increases the possibility of self-pollen deposition, the effect of which depends on the mating system. The cost of selfpollen deposition is through inbreeding depression (reduced fitness of offspring produced by selfing) in self-compatible species (Charlesworth and Charlesworth 1987, Husband and Schemske 1996) or through sexual interference (Lloyd and Webb 1986, Webb and Lloyd 1986, Bertin and Newman 1993, Barrett 2002), which leads to wastage of gametes in both self-compatible and self-incompatible species (Lloyd and Webb 1986, Webb and Lloyd 1986, Fetscher 2001, Barrett 2002). Selection to reduce self-pollen deposition appears to have played a significant role in floral evolution (Lloyd and Webb 1986, Webb and Lloyd 1986). Two broad strategies have evolved – spatial separation (herkogamy) or temporal separation (dichogamy) of sporophylls (i.e., the male and female reproductive parts) (Armbruster et al. 2009). These strategies lead to a further complication in animal-pollinated plants. For maximum pollination accuracy, pollen must be released and received from the same position (called the presentation zone), because animal pollinators gather and dispense pollen from the same part of their body. Therefore, in a dichogamous species, the first sporophyll has to be removed from the presentation zone so that the second sporophyll can take its place (Lloyd and Webb 1986, Medan and Ponessa 2003). This can be achieved in three ways – by abscission, movement, or shrivelling of the first sporophyll. In this manner, anthers and stigma are positioned in the presentation zone, in any sequence, with a temporal separation of male and female phases. In protandrous species, anthers are positioned first at the presentation zone during the male phase, and the stigma is presented later in the female phase; this sequence is revered in protogynous species. Dichogamy is widespread among hermaphroditic flowers (Bertin and Newman 1993). However, only a few of these species show movement-assisted dichogamy – where sporophylls move in a direction opposite to each other: the first sporophyll is removed by movement and then the second takes its place (Medan and Ponessa 2003, Ruan and da Silva 2011).

Early naturalists considered dichogamy as a perfect adaptation, where the second sporophyll takes precisely the place of the first one (Gray 1879, Kerner von Marilaun and Oliver 1895, Knuth and Müller 1908). The proposition of perfection is based on three assumptions. First, the positional exchange of sporophylls is precise, such that anther and stigma touch the same body part of the pollinator (Armbruster et al. 2009). At the population level, this increases pollination accuracy and thus also increases pollen delivery to conspecific stigmas. Second, the position that both sporophylls take is optimal, in that they touch that body part of the pollinator which results in maximum pollen transfer (Armbruster et al. 2009). Third, the position of sporophylls should be stable in the presentation zone, hence maximising their duration at this position, which should result in the maximum transfer of pollen.

In movement-assisted dichogamy, movements of sporophylls in opposite directions separate male and female phases temporally. However, the possibility of self-pollen deposition cannot be altogether avoided because in the transition phase, when sporophylls cross each other, self-pollen can still be deposited as the stigma-anther distance is minimal at that time. Although this transition phase is brief, one would expect floral adaptations to minimise this micro-scale problem inherent in movement-assisted dichogamy. This problem could potentially be minimised in two ways – either by minimising the deposition of self-pollen, or if deposited, by minimising the chance of fertilization. In protandrous species, the deposition of self-pollen during the crossover period can be reduced by delaying the opening of the stigma lobes until after the sporophylls have crossed. This is a physical solution to the problem: a closed or narrowly opened stigma lobe reduces the available stigma surface area, and makes pollen deposition difficult despite the closeness of stamen and style. A second possible solution is chemical: if stigma receptivity is delayed until later, any pollen deposited in the transition phase would not adhere to the surface or germinate. This hypothesis predicts that the proportion of receptive stigmas are higher in the later part of female phase than the earlier part of the female phase.

To investigate these phenomena we studied *Clerodendrum infortunatum* Gaertn. [Lamiales: Lamiaceae] (henceforth *Clerodendrum*) flowers, which exhibit movement-assisted protandrous dichogamy. We examined the coordination of sporophyll movements, measured the stigma lobe angle as indicator of stigma lobe openness, and counted the number of deposited pollen grains on stigmas collected at different floral phases. One would expect that stigma lobes should be fully open in the female phase, and stigmas should become receptive only at that phase; the consequence should be higher pollen deposition in the female phase than the male phase.

## Methods

The Western Ghats mountain range stretches along the west coast of India. The Anamalai Hills, dominated by mid-elevation tropical wet evergreen rainforest (Pascal 1988), is located in the southern Western Ghats. The Valparai plateau (700-1600m asl), which is dominated by tea and coffee plantations, lies within these hills. However, many private rainforest fragments persist in the landscape. These scattered, disturbed forest patches harbour a significant proportion of rainforest flora and fauna (Umapathy and Kumar 2000, Sridhar et al. 2008). Two such forest fragments were selected for this study, namely Iyerpadi Riverine (10°21’12” N, 76°59’54” E, Area 1.76 ha) and Varattuparai Colony (10°21’22” N, 76°56’30” E, Area 33.4 ha).

Within these forest fragments, we studied *Clerodendrum*, a native shrub, between December 2013 and May 2015. *Clerodendrum* has a long flowering season, which extends from December to May (Supplementary Fig. 1). It produces white flowers, which are mostly visited by butterflies (Reddy et al. 1995, Kumar et al. 2017). The flowers are hermaphrodite and complete, their five petals forming a tubular corolla 1.5 cm long, which opens out at the distal end. Four long stamens and one style, all of approximately the same length protrude from the corolla (Kumar et al. 2017).

Sporophyll movements were monitored throughout the floral lifetime of a total of 32 flowers from 18 plants. Flower monitoring was carried out between 18 February 2015 and 26 April 2015. Buds were tagged with lengths of coloured thread and were monitored every 2 hours after anthesis. Monitoring began at 06:00 AM, with seven observations per day until nightfall, and then continuing the next morning. We used hr 2 anther data as the initial anther stage because at hr 0 (at anthesis) a large portion of the anthers remains undehisced, and stamens remain coiled (personal observation). During each observation, a photograph of the lateral view of the flower was taken, and the distance from the style to each of the four anthers was measured with vernier callipers to the nearest 0.01 mm. Stamen and style angles were measured from those photographs using ImageJ software, Fiji distribution (Schindelin et al. 2012), in an anti-clockwise direction from the basal end of the flower along the floral axis, which is approximately horizontal (see Fig. 1 for an illustration). Stigma lobe angles were also measured. To assign phases to flowers, we used the position of the sporophylls together with the angle between the stigma lobes. Movement data show that the crossing of male and female sporophylls occurs at around 180° angle (Fig. 2). From this, we assigned the flower to be in the male phase when the average anther angle was below 180° (i.e., above the floral axis). If the average anther angle was above 180° (i.e., below the floral axis), we labelled the flower to be in a ‘neutral’ phase if the stigma lobe angle was less than 70° (such that the probability of the stigma being receptive was less than 0.5, see Results) or in the female phase if the stigma lobe angle was greater than 70°. Note that these phases were defined only with respect to the position and the shape of the sporophylls. Phase durations were calculated with a precision of 2 hrs for the male and neutral phase and with a precision of 12 hrs for the female phase.

**Fig. 1.**
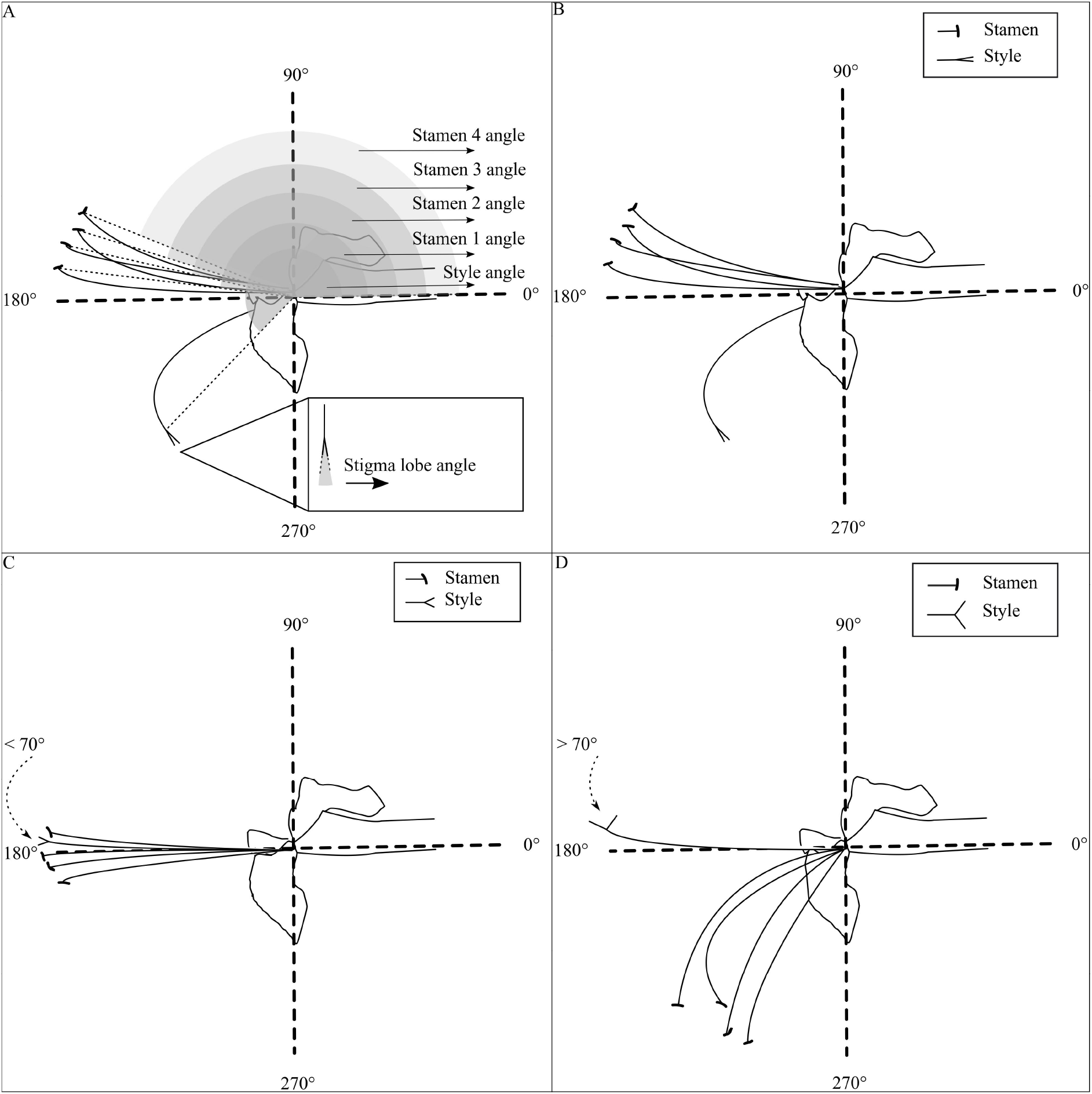
Schematic diagram of *Clerodendrum* flower (lateral view). The diagram is not to scale. For clarity, only three of five petals are shown. The dotted lines are floral axes. A. Schematic of angle measurement. B. Male phase flower C. Neutral phase flower D. Female phase flower.

**Fig. 2.**
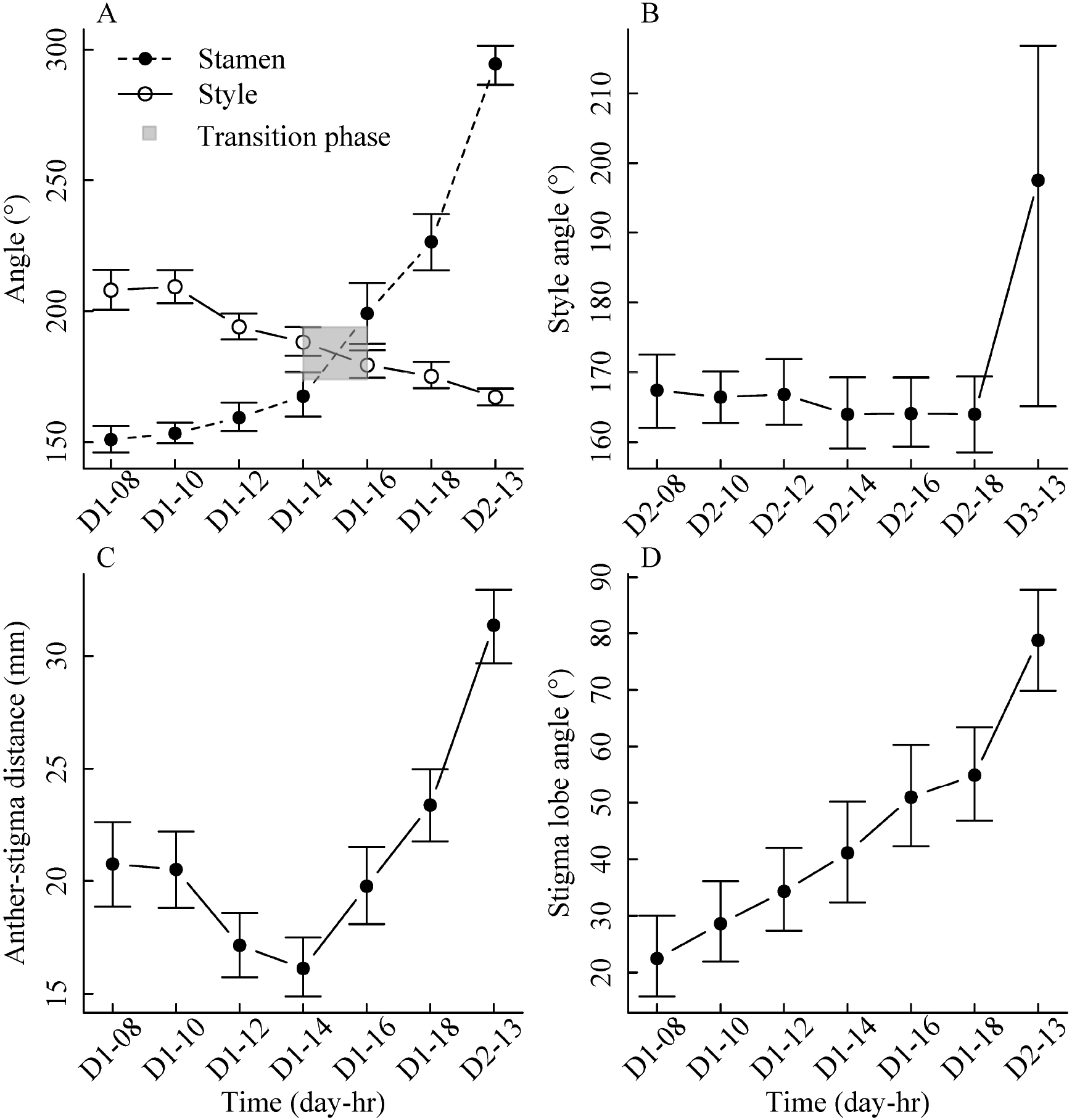
Movement dynamics of sporophylls in the floral lifetime of *Clerodendrum*. Time on the X-axis is denoted according to day and 24-hour clock format. (A) Stamen and style angle (B) Style angle on the 2^nd^ day (C) Anther-stigma distance (D) Stigma lobe angle. Error bars are 95% nonparametric bootstrap confidence intervals (N = 32 flowers). In panel A and B, increasing angle indicates dropping of the sporophyll.

Stigma receptivity was tested in 24 stigma samples collected from different flower stages in the field on 28 May 2015. This was done using a DAB (Sigma Fast™ 3.3’ Diaminobenzidine, SIGMA D-4168) reaction test, which tests for the presence of peroxidases on the receptive stigma surface (Rodriguez-riano and Dafni 2000, Dafni et al. 2005). High peroxidase enzyme activity is related to stigmatic receptivity (Dafni et al. 2005). Excised stigmas were immediately placed on separate slides, drops of reagents added, and samples covered with DPX mountant (Sigma Aldrich) and coverslips. The stigma samples were brought back in the lab within 24 hr and observed under the microscope for the presence of stain. If the stain was present in the stigma sample, the stigma was considered to be receptive. Also, photographs were taken of the stigma lobes (at an angle orthogonal to the lobes) and angles between the lobes were measured using ImageJ.

To estimate the stigmatic pollen load at different floral phases, stigmas were collected haphazardly from 18 plants during different floral phases (21 stigmas in male phase, 23 in neutral, 24 in female) between 07 April 2015 and 14 April 2015. Collected stigmas were immediately placed in separate vials containing 70% ethanol for lab analysis. This storage method has a caveat – pollen grains loosely attached to the stigma surface may get washed off. Still, pollen that remain adhered on the stigma following this preservation technique are likely to give a reasonable estimate of the number of germinating pollen on stigmas (Vazquez and Simberloff 2004, Ortega et al. 2007). In the laboratory, pollen counting was done between 21 June 2016 to 30 June 2016. Stigmas were excised, placed on microscope slides, washed with 95% ethanol and stained with Safranin O (Green 1990, Wood et al. 1996, Jones 2012). Glycerine was added to the stigmas, and they were squashed under a coverslip. The number of pollen grains were counted under a compound microscope (100x magnification).

All analyses were carried out in R, version 3.5.1 (R Core Team 2018); together with the additional packages: nlme (Pinheiro et al. 2018) and boot (Davison and Hinkley 1997, Canty and Ripley 2017). Error bars in all figures are 95% nonparametric bootstrap confidence limits. The nonparametric bootstrap method circumvents restrictive assumptions about the underlying distribution from which the data were drawn (Chernick and LaBudde 2014). We draw inferences from the data based on the estimated effects and their uncertainty. In addition to estimating effect sizes (Nakagawa and Cuthill 2007, Wasserstein and Lazar 2016, Wasserstein et al. 2019), we also present results of null hypothesis significance tests.

## Results

Anthesis happens in the morning. After that, stamens and style show broadly opposite movement directions over time (Fig. 2). Starting from around 150° at hr 2, stamens gradually fall to the floral axis and then below. At hr 2, the style is positioned at around 210°, remaining roughly stable at that position (with a slight downward movement) before beginning a gradual but steady rise at hr 4. Stamens and style cross each other at about 8 hrs (transition phase) after anthesis (Fig. 2A). The next day the style reaches around 165° and remains there for an entire day (Fig. 2B). However, for the stamens, there is no similar stable position when they remain in the presentation zone; instead they keep moving. At the transition phase, the anther-stigma distance is minimal (Fig. 2C). The angle between the stigma lobes is at about 20° at hr 2, and the lobes progressively open throughout the floral lifetime. However, during the transition phase stigma lobes remain only narrowly opened (around 40°) (Fig. 2D).

Style and average stamen angle are negatively correlated (Spearman’s rank correlation: S = 2497000, p < 0.001, ρ = −0.64, Fig. 3). Reading Figure 3 in temporal sequence from right to left illustrates that the initial large change in style angle is accompanied by only a small change in average stamen angle. Later in the sequence, however, a small change in style angle corresponds with a high increase in (i.e., lowering of) average stamen angle.

**Fig. 3.**
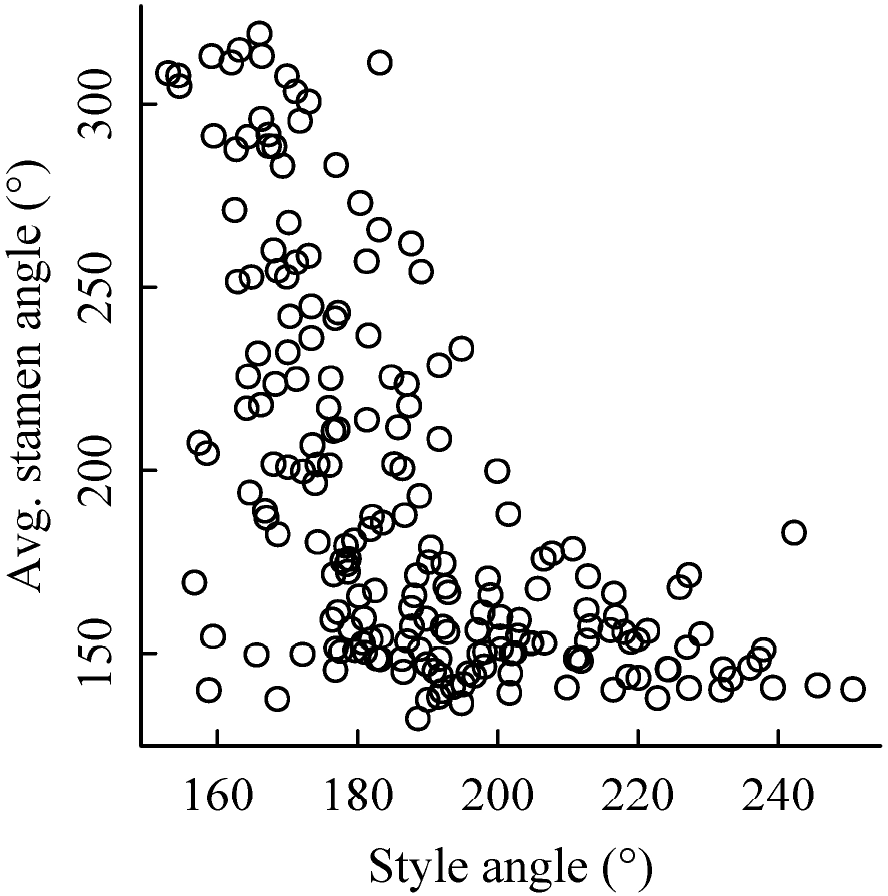
Relationship between the style angle and average stamen angle in *Clerodendrum* floral lifetime

Stigma receptivity tests show that stigmas are not receptive during the male phase. However, not all stigmas are receptive in the female phase either. Only around 55% of the stigma samples were judged receptive (by the method used) in the female phase. Stigma receptivity was initially modelled as a binomial outcome (receptive vs non-receptive) in a generalised linear model (GLM) with two predictors – style angle and stigma lobe angle. However, these two predictors were highly correlated with each other and so we present results for a model with stigma lobe angle (which explained more variability) as the sole predictor (McFadden pseudo-*R^2^* = 0.57; likelihood ratio test: *χ^2^* (1, *N* = 24) = 4.40, *p*<.05). A 1° increase in stigma lobe angle, is associated with an increase in the odds of the stigma being receptive (versus not being receptive) by a factor of 1.09. The plot suggests that the probability of stigma receptivity is 50% at roughly 70°: we define this angle as the critical stigma lobe angle (Fig. 4).

**Fig. 4.**
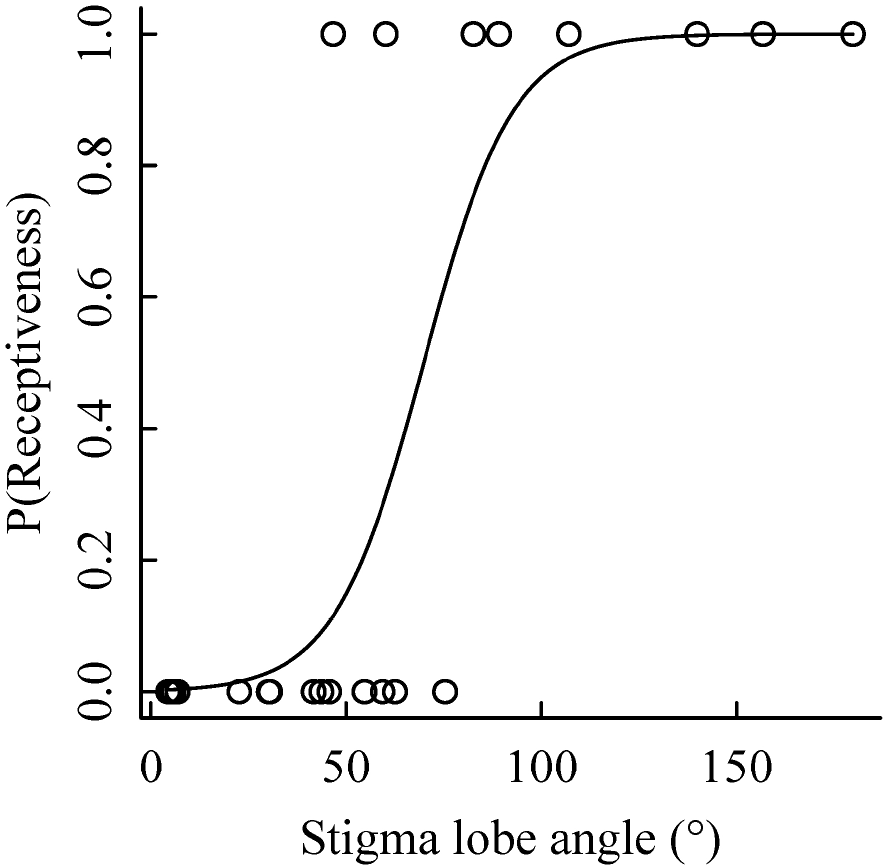
A generalised linear model is fitted to the data displaying the relationship between stigma lobe angle and stigma receptivity (N = 24).

Solely based on stamen and style movement, *Clerodendrum* flower has two distinct phases – male phase (average anther angle below 180°) and the female phase (average anther angle above 180°), assuming that floral phases do not change at night (Fig. 2). However, based on the stigma lobe angle and stigma receptivity, *Clerodendrum* flowers have three distinct phases. The male phase lasts for an average of 7.8 hrs [95% nonparametric bootstrap CI: 7.0, 8.7; *N* = 28 flowers], whereas the female phase lasts for an average of 40.8 hrs [95% nonparametric bootstrap CI: 36.0, 45.9; *N* = 28 flowers]. There is an additional neutral phase between them, which lasts for an average of 3.6 hrs [95% nonparametric bootstrap CI: 2.5, 4.5; *N* = 28 flowers]. Just after the male phase is over, although the average anther angle is above 180° (as the transition takes place at 180°, so the style is below 180°), the stigma is not yet receptive (Fig. 4) and stigma lobes are not fully opened (Fig. 2). So we conclude that this short phase is neither in the male nor the female phase and we refer to it as the ‘neutral’ phase.

No pollen was found on male-phase stigmas, and while a few occurred on neutralphase stigmas, the maximum pollen deposition occurred on female-phase stigmas (Fig. 5). A one way ANOVA showed that the differences in pollen deposition between the male (*N* = 21, *M* = 0, *SD* = 0), neutral (*N* = 23, *M* = 0.47, *SD* = 0.99) and the female phase (*N* = 24, *M* = 1.91, *SD* = 3.17) were statistically discernible (*F*(2,65) = 5.851, *p*< 0.01). Planned contrast revealed that neutral stage pollen deposition is statistically discernibly higher than male stage pollen deposition, *t* = −2.31, *p*< 0.05. Also, female stage pollen deposition is discernibly higher than neutral stage pollen deposition, *t* = −3.34, *p*< 0.01.

**Fig. 5.**
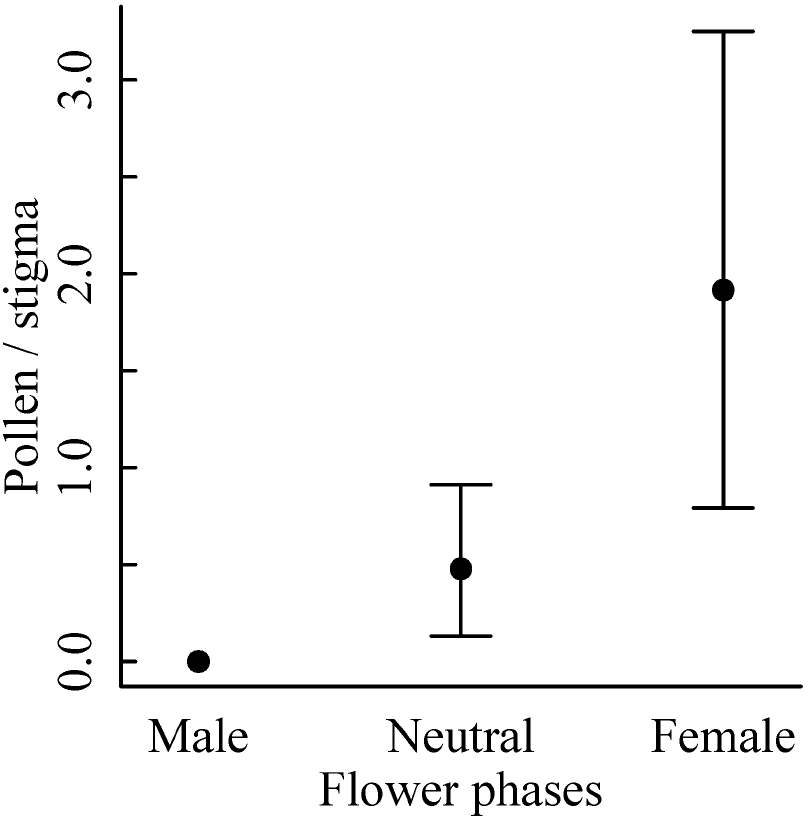
*Clerodendrum* pollen deposition on male (N = 21), neutral (N = 23) and female phase (N = 24) stigmas. Error bars represent 95% nonparametric bootstrap confidence intervals.

## Discussion

This study illuminates the details of sporophyll dynamics in *Clerodendrum*. The species illustrates a typical case of movement-assisted dichogamy, where the bending ability of the sporophylls minimizes self-pollen deposition (Medan and Ponessa 2003, Ruan and Silva 2011). Flowers have three distinct phases: male, female and an in-between ‘neutral’ phase. The female phase is much longer than the male phase, and the neutral phase is the shortest in duration. The male phase precedes the female phase, with the transition between the phases taking place around eight hours after anthesis (Fig. 2A). During the time of transition, flowers pass through the neutral phase, when the stigma-anther distance becomes minimal, making self-pollen deposition likely (Fig. 2C). However, at this point, the stigma lobe angle remains narrow (Fig. 2D), and the stigma remains non-receptive as stigma receptivity is closely related to stigma lobe angle (Fig. 4). Hence both physical and chemical control of pollination occurs in synchrony to minimise the chance of self-pollen deposition.

How do these findings relate to the assumption that sporophylls are stable in the presentation zone? The movement dynamics show that the style remains nearly stationary throughout the female phase (Fig. 2B). By contrast, stamens are in constant motion during the male phase, and therefore they show no single stable position. As a result of this, there is a wide range of stamen angles from which pollen export takes place. This wide range may serve to ensure pollen deposition in a population where variance of style angle is high. However, there is a price to be paid in terms of inaccurate pollen export, where some pollen is exported from stamen angles which are away from the mean style angle in the population, therefore resulting in lower pollination accuracy. In this way there is a trade-off between reproductive assurance and pollination accuracy. Such a trade-off is likely to occur in other dichogamous species as well. In future studies, the consequences of dynamic anther position needs to be explored in more detail.

Early pollination biologists were aware of sporophyll movement in flowers (Lindley 1819, 1826). Sporophylls usually move in a combined and coordinated manner (Lloyd and Webb 1986, Bertin and Newman 1993, Ruan and Silva 2011). The adaptive significance of sporophyll movements includes reducing selfing, increasing outcrossing, reducing sexual interference and helping in delayed selfing in a harsh environment (Ruan and Silva 2011). As *Clerodendrum* is a self-compatible species and breeds by mainly xenogamy and geitonogamy (Reddy et al. 1995, Kumar et al. 2017), the possible ways sporophyll movement can benefit the plant are by reducing both selfing and sexual interference. In *Clerodendrum*, the style and stamen move in a direction opposite to each other, and they do so in a coordinated manner. In general, correlations among floral traits are higher than correlations among vegetative traits in plants because flowers are under direct selection (Berg 1959, Hansen et al. 2007). As expected, the correlation between the positions of style and stamens at different time points in a floral lifetime is high (Fig. 3). Such correlations could be the result of pleiotropy or correlated selection. Correlated selection occurs when a specific combination of trait values have higher fitness than other combinations (Cheverud 1982, Lande and Arnold 1983, McGlothlin et al. 2005). Traits that are functionally related are more likely to be under correlated selection. Thus the relationship between style and stamen movement in *Clerodendrum* could be under correlational selection. In future, the cause behind the correlation between style and stamen movement in *Clerodendrum* needs to be studied.

Another noteworthy characteristic is the shape of the correlation. Style and stamens at different time points are non-linearly correlated. The adaptive significance of the shape of this relationship also needs to be studied in detail.

The sporophyll movements give rise to two functional floral phases which are temporarily separated. The male phase, which comes first, is much shorter in comparison with the later female phase. During the phase transition in movement-assisted dichogamous plants, such as *Clerodendrum*, a flower faces a unique situation. The crossing of sporophylls creates the very problem which it tries to solve – the problem of self-pollen deposition – as the distance between anther and stigma is minimal at this point (Fig. 2C).

An intermediate neutral phase is a plausible solution to this problem, in which the effect of self-pollen deposition is minimised by keeping male and female phases separated. Complete dichogamy occurs when male and female phases are separated and without any overlap. On the other hand, incomplete dichogamy occurs when there is an overlap. *Clerodendrum* shows complete dichogamy as there is no overlap between male and female phase because of the neutral phase that exists in between, as previously argued. It is expected as complete dichogamy should be accompanied by a neutral phase (Aizen and Basilio 1995). A combination of ecological factors, including pollen limitation and inbreeding depression, influences whether complete or incomplete dichogamy is the best strategy for the plant (Rosas-Guerrero et al. 2017). In future, the link between pollen limitation and complete dichogamy in *Clerodendrum* needs to be rigorously tested.

The neutral phase in *Clerodendrum* is created by the combined actions of delayed stigma lobe opening and delayed stigma receptivity. Delayed stigma lobe opening reduces the active stigma surface area that might come in contact with self-pollen. Delayed receptivity prevents self-pollen from blocking the stigma surface. A receptive stigma is sticky and therefore binds pollen grains more efficiently than a non-receptive stigma, from which pollen grains are easily dislodged. For this reason, self-pollen have a lower chance of germinating and blocking the ovule when stigmas are not receptive. Our results suggest that stigma receptivity is closely associated with stigma lobe angle (Fig. 4). The stigma lobe angle is quite narrow (35°) in the transition phase (Fig. 2D). Thus during the transition phase stigmas are non-receptive in *Clerodendrum*. Indirect support for these results come from another study on *Clerodendrum* that carried out hand pollination during different times in floral development (Reddy et al. 1995). The resultant seed set showed that the stigma becomes receptive 12 hours after anthesis. In our study, we show that the transition phase happens around eight hours after anthesis (Fig. 2A). Taken together, stigma lobe angle, chemical receptivity and seed set suggest that the flower does not yet enter the female phase during transition. Although it appears that the phase transition occurs when the stamens and style cross each other, in reality, style and stamen angles are poor indicators of phase transition (Fig. 4). Flower enters the female phase only when the stigma becomes receptive, irrespective of style angle. When stamens and style cross each other, the male phase is usually over (depending on pollen availability on the anther, which indirectly depends on visitation rate) and the flower enters into a brief neutral phase before proceeding to the female-phase, thus minimising the problem of self-pollen deposition.

If these two mechanisms work well enough to solve the problem of self-pollen deposition in the transition phase, pollen deposition should be minimal in the neutral phase. This prediction turns out to be broadly true, with pollen deposition in neutral phase being considerably less than that in the female phase (Fig. 5). Thus the existence of a neutral phase resolves the problem of self-pollen deposition to a great extent in *Clerodendrum*. Such an adaptation may be quite common: a study on *Alstroemeria aurea* also describes how a neutral phase had resolved the conflict (Aizen and Basilio 1995).

The elaborate movement mechanisms at the level of a flower might still fail to achieve the plant level goal (de Jong et al. 1992, Galloway et al. 2002, Duan et al. 2005). Geitonogamous self-pollen deposition across flowers from the same plant might reverse the reduction in self-pollen deposition that dichogamy enables at the flower level (i.e., pollen deposition from the anthers of one flower to its own stigma). Dichogamy in *Clerodendrum* is asynchronous, and male- and female-phase flowers are found together at the same time and on the same plant. So potentially, geitonogamy can take place in *Clerodendrum*, and indeed it does (Reddy et al. 1995, Kumar et al. 2017). In general, geitonogamy is detrimental to plants (de Jong et al. 1992), therefore any mechanism to reduce it would be advantageous. Lowering the number of open flowers could be an effective strategy as plants with larger floral display show higher geitonogamy (de Jong et al. 1992). In *Clerodendrum*, less than 20% of buds are open even at the peak of the flowering season (Supplementary Method, Supplementary Fig. 1), which could effectively reduce the extent of geitonogamy. Overall it seems that multiple plant traits apart from dichogamy, such as low open flower to bud ratio and extended blooming, might play a role in reducing overall self-pollen deposition in *Clerodendrum*. Further study is needed to tease apart their relative contributions.

In general, the floral adaptations of movement-assisted dichogamous species must be fine-tuned to handle the problem of self-pollen deposition successfully and to maximise the reproductive fitness in hermaphrodite flowers. Several accessory adaptations ranging from physical to chemical may work in tandem to achieve this. In *Clerodendrum*, delayed stigma lobe opening, together with delayed stigma receptivity work together to solve the transition phase problem.

