## Supplementary Information for "Fine-tuned spatiotemporal dynamics of sporophylls in movement-assisted dichogamy: a study on *Clerodendrum infortunatum*"

The following Supplementary Information is available for this article:

**Supplementary Method** Overall flowering phenology can be measured at two levels: at the level of the individual plant, to assess how the number of flowers per plant changes over time; and at the level of the population, to measure how the proportion of plants in flower changes with time. To study plant-level flowering phenology, 28 individual plants were monitored every 15 days from 13 January to 16 May 2014. At each visit, the total number of individual open flowers and buds (across all inflorescences) were counted on each plant. *Clerodendrum* is an edge species, and to understand its degree of habitat specialization, belt transects (3m wide and 50 m long) were laid from the edge to the interior of forest fragments to measure population-level phenology. Twenty-three transects were laid in fragments locations where *Clerodendrum* was present, with a minimum distance of 30m between two adjacent transects. In every 5m segment of each transect, the number of *Clerodendrum* plants taller than 1m was counted. On each plant, I noted the presence or absence of flowers (including buds). These surveys were carried out once a month from 23 December 2013 to 15 April 2014.

**Supplementary Fig. 1 Flowering phenology of *Clerodendrum* *infortunatum. Clerodendrum* has an extended flowering season, which stretches at least from December to May. (A) At the level of the individual plant (n = 28 plants), the percentage of open flowers increases from January, peaks in end February, and declines thereafter. However, fewer than 20% of buds are open at any given time. (B) At the population level (n = 23 transects), the percentage of flowering individuals follows a similar trend, peaking in February-March. Even at the peak, however, fewer than 40% of the plants are in flower. Error bars are bootstrapped 95% nonparametric confidence intervals.**

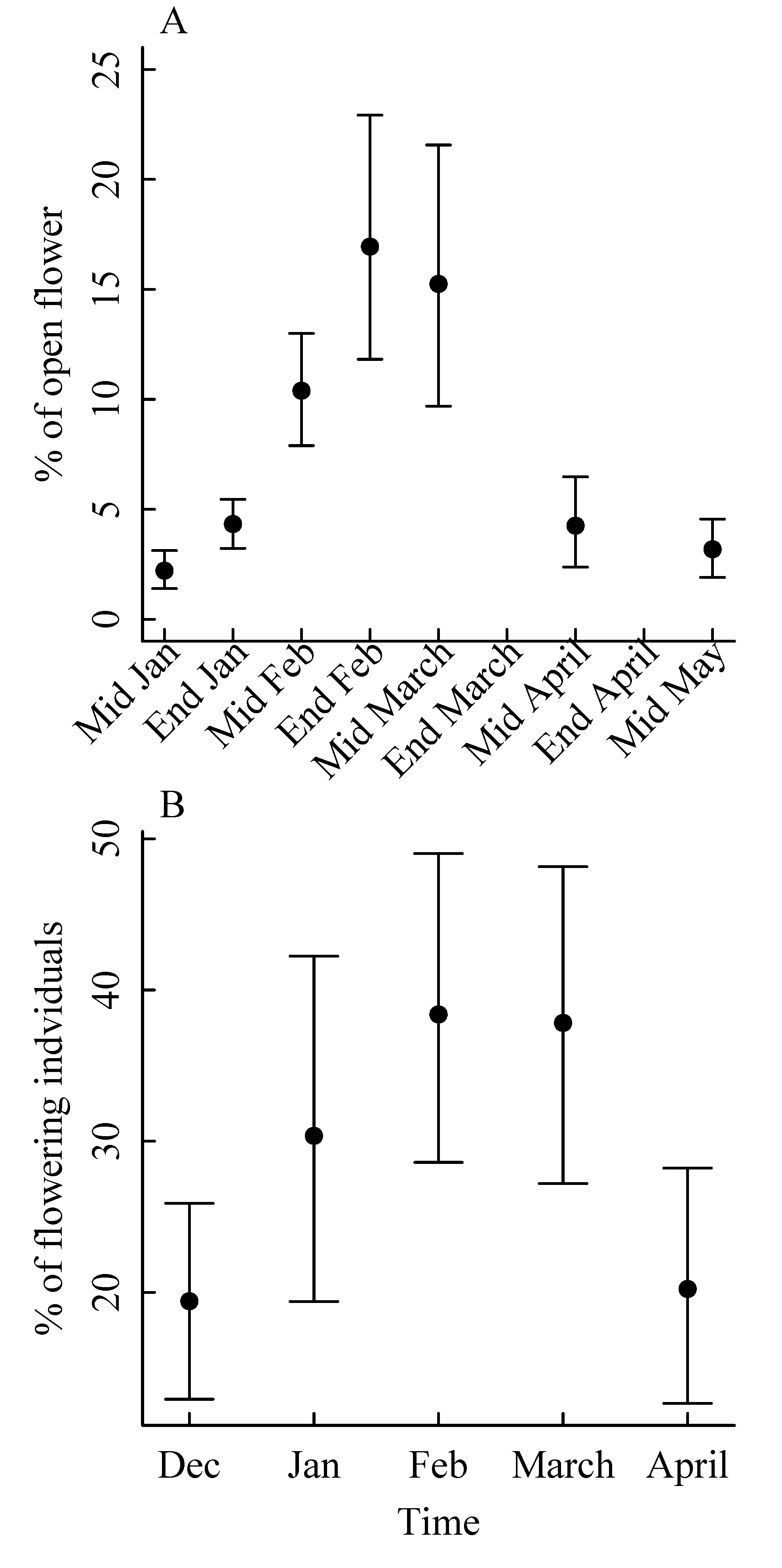
